# Motor learning selectively strengthens cortical and striatal synapses of motor engram neurons

**DOI:** 10.1101/2021.10.28.466357

**Authors:** Fuu-Jiun Hwang, Richard H. Roth, Yu-Wei Wu, Yue Sun, Yu Liu, Jun B. Ding

**Affiliations:** Department of Neurosurgery, Stanford University, Stanford, CA 94305, USA; Institute of Molecular Biology, Academia Sinica, Nankang, Taipei, Taiwan; Department of Neurology and Neurological Sciences, Stanford University, Stanford, CA 94305, USA

**Keywords:** motor learning, corticostriatal circuit, memory engram, synaptic plasticity, dendritic spines, synaptic clustering, two-photon imaging

## Abstract

Learning and consolidation of new motor skills require adaptations of neuronal activity and connectivity in the motor cortex and striatum, two key motor regions of the brain. Yet, how neurons undergo synaptic changes and become recruited during motor learning to form a memory engram remains an open question. Here, we train mice on a single-pellet reaching motor learning task and use a genetic approach to identify and manipulate behavior-relevant neurons selectively in the primary motor cortex (M1). We find that the degree of reactivation of M1 engram neurons correlates strongly with motor performance. We further demonstrate that learning-induced dendritic spine reorganization specifically occurs in these M1 engram neurons. In addition, we find that motor learning leads to an increase in the number and strength of outputs from M1 engram neurons onto striatal spiny projection neurons (SPNs) and that these synapses form local clusters along SPN dendrites. These results identify a highly specific synaptic plasticity during the formation of long-lasting motor memory traces in the corticostriatal circuit.

**HIGHLIGHTS:**

– Motor performance is correlated with the reactivation of motor engram neurons
– Motor learning increases spine density and new spine survival selectively on M1 engram neurons
– Motor learning strengthens motor engram outputs to the striatum
– M1 engram outputs converge onto clusters of dendritic spines on striatal spiny projection neurons

## INTRODUCTION

Learning and executing new motor skills are crucial functions of the brain and involve the coordinated activity of the motor cortex and basal ganglia. Notably, the activity patterns of neurons in the primary motor cortex (M1) as well as of spiny projection neurons (SPNs) in the striatum adapt over the course of motor learning and becomes more closely associated with learned movements (Jin and Costa, 2010; Peters et al., 2014; Barbera et al., 2016; Klaus et al., 2017; Sheng et al., 2019). An intriguing interpretation of these adaptations in neuronal activity is that such behavior-related neurons may represent the neural correlate of motor memory, forming a motor memory engram.

Memory engram neurons have been identified in a variety of brain regions and have been associated with a wide spectrum of behaviors, ranging from associative learning to aversive and reinforcing memories or maladaptive behaviors (Liu et al., 2012; Hsiang et al., 2014; Ramirez et al., 2015; Girasole et al., 2018; DeNardo et al., 2019; Josselyn and Tonegawa, 2020). Recently developed genetic tools that use immediate-early gene promoters to label neurons that are active during specific stages of learning have allowed for the characterization of these engram neurons and for defining their roles during behavior (Reijmers et al., 2007; Guenthner et al., 2013; Denny et al., 2014; DeNardo et al., 2019). A long-standing question in the field is what cellular and synaptic processes drive the formation of engram neurons during learning and how these neurons become integrated into brain circuits.

In the motor cortex, synaptic plasticity on dendrites of Layer 5 pyramidal neurons (L5PNs) play a crucial role in the acquisition of new motor skills (Rioult-Pedotti et al., 2000; Xu et al., 2009; Yang et al., 2009; Guo et al., 2015; Roth et al., 2020; Albarran et al., 2021). Specifically, motor learning induces enhanced formation of new dendritic spines and a delayed increase in spine elimination, which results in a transient increase in spine density on L5PNs. The rate of spine formation and especially the stabilization of newly formed spines correlates with motor performance (Xu et al., 2009; Albarran et al., 2021). Furthermore, artificially eliminating learning-induced new spines impairs learned motor behaviors (Hayashi-Takagi et al., 2015). While these experiments revealed a motor memory trace at the inputs of M1 neurons, very little is known about how this specific synaptic mechanism is related with motor engram neuron activity patterns, and in addition, how the outputs of these neurons change during learning.

A major target of M1 output neurons is the dorsolateral striatum (DLS), where they form glutamatergic synapses onto postsynaptic striatal SPNs. This corticostriatal projection is crucially involved in motor learning, as highlighted by loss-of-function studies. DLS lesions or silencing SPNs impairs learned motor behaviors, and blocking SPN plasticity by deleting NMDA receptors on SPNs prevents mice from learning new motor skills (Dang et al., 2006; Santos et al., 2015; Sheng et al., 2019). Importantly, in psychogenic movement disorders, such as Parkinson’s disease, autism spectrum disorders, and L-DOPA-induced dyskinesia, disruption of ensemble activity of neurons in the DLS or M1 may mediate behavioral deficits (Parker et al., 2018; Maltese et al., 2021; Yin et al., 2021). Conversely, neuronal engrams that are not present in the normal brain are formed in the striatum during maladaptive motor behaviors (Girasole et al., 2018). Finally, loss of dopamine also leads to aberrant spine plasticity in M1 and striatal SPNs (Day et al., 2006; Guo et al., 2015; Xu et al., 2017).

The widespread and convergent innervation of corticostriatal projections has made it challenging to assess the function and plasticity of this circuit over the course of motor learning. To study how the corticostriatal circuit adapts during motor learning, we used the genetic Targeted Recombination in Active Populations (TRAP) (Guenthner et al., 2013; DeNardo et al., 2019) approach to identify and label behavior-relevant engram neurons in M1 and examined their input and output synaptic plasticity in mice learning a forelimb reaching task. We found that a population of behavior-related engram neurons emerged in M1 during motor learning in well-trained learner mice. Surprisingly, we found that motor learning-induced a transient increase in dendritic spine density and stabilization of newly formed spines specifically on engram neurons but not neighboring non-TRAPed neurons. We further found that motor learning led to a selective strengthening of M1 engram neuron outputs formed onto the clustered spines of postsynaptic SPN dendrites. Our study reveals a highly selective synaptic plasticity mechanism in corticostriatal circuits that lead to the formation of long-lasting motor memories.

## RESULTS

### M1 engram neurons are reactivated when performing a learned motor task

One of the hallmarks of memory engrams is that an engram formed during learning is subsequently reactivated during memory retrieval (Semon, 1904; Josselyn and Tonegawa, 2020). In M1, cortical neurons form ensembles, and their firing patterns during motion sequences encode task-relevant features (Costa et al., 2004; Ölveczky et al., 2005; Jin and Costa, 2010). However, whether these neurons that are activated during motor learning are reactivated during the performance of a previously learned motor task has not been tested. To label motor behavior-activated neurons in M1 during motor learning, we trained c-fos TRAP mice to perform a forelimb reaching task, in which mice use a single paw to reach for a small food pellet through a slit opening in the training chamber (Figure 1A) (Xu et al., 2009; Guo et al., 2015; Roth et al., 2020; Albarran et al., 2021). Over the course of eight days of daily training, mice increased their rate of successfully retrieving the pellet (Figure 1B). Using the c-fos TRAP system, neurons active during a specific time window can be labeled with the fluorescent protein tdTomato by injecting 4-Hydroxytamoxifen (4-OHT) in c-fos-CreER mice crossed to a tdTomato reporter mouse (Ai9). By injecting 4-OHT either during the first two days of training (early TRAP) or during the last two days of training (late TRAP), we specifically labeled cells activated during the early or late stages of motor learning. A week after TRAP labeling, mice were trained for one additional session and perfused within an hour after training to allow for quantification of cells activated during the latest session by c-fos immunostaining. In late TRAP mice, this approach allows us to identify which neurons were activated during late-stage training - when mice have learned to perform the reaching task (TRAP neurons) - and which neurons were activated or reactivated in the training session one week later when mice still successfully perform the task (c-fos positive neurons) (Figure 1C and 1D). Early TRAP mice were only trained for two days with a third training session a week after initial training, allowing us to measure neuron activation and reactivation during the early stages of learning when the mice have not yet acquired the motor skills. Control mice were not trained but exposed to the training chamber and given 4-OHT in parallel with the early or late TRAP mice.

**Figure 1:**
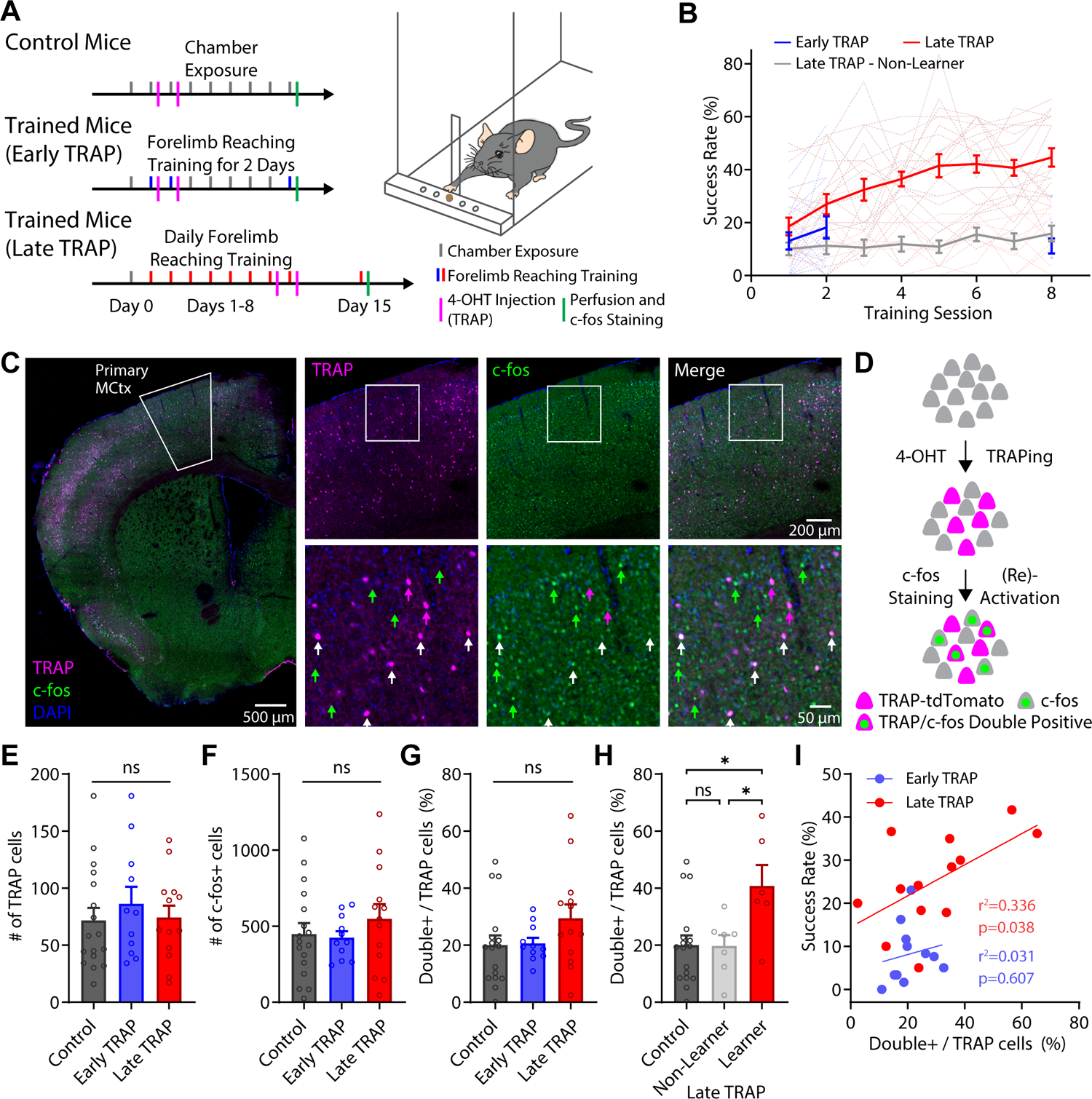
Reactivation of motor cortex engram neurons correlates with behavioral performance on the forelimb reaching task. (A) Experimental timeline and schematic drawing of the reaching task. (B) Behavioral performance of mice trained on the forelimb reaching task. Thin lines represent individual mice, and the bold line is the average. n = 17 Early TRAP mice, 21 Late TRAP Learner mice, 10 Late TRAP Non-Learner mice. Error bars, SEM. (C) Representative image of TRAP-tdTomato (magenta) fluorescence and c-fos immunostaining (green) in the primary motor cortex (M1). Left: an overview of entire brain hemisphere, upper right: enlarged view of M1, lower right: enlarged view of the white square in M1. Green arrows denote cells with only c-fos expression, magenta arrows denote cells with only TRAP labeling, white arrows denote cells with c-fos expression and TRAP labeling. (D) Schematic drawing showing TRAPing of activated neurons with 4-OHT in c-fos TRAP-Ai9 mice (magenta) and immunostaining for c-fos proteins (green). Reactivated neurons show TRAP/c-fos double-positive signals. (E) Number of TRAP-tdTomato labeled cells in M1 (analyzed area: ∼1.13 mm^2^). Bars denote mean and circles individual mice. Error bars, SEM. Control: 71.77 ± 10.93, n = 17 mice; Early TRAP: 86.34 ± 14.81, n = 11 mice; Late TRAP: 74.19 ± 10.64, n = 13 mice. p = 0.681, One-way ANOVA. (F) Number of c-fos-expressing cells in M1 (analyzed area: ∼1.13 mm^2^). Bars denote mean and circles individual mice. Error bars, SEM. Control: 447.4 ± 72.36, n = 17 mice; Early TRAP: 424.3 ± 42.92, n = 11 mice; Late TRAP: 548.1 ± 96.92, n = 13 mice. p = 0.513, One-way ANOVA. (G) Fraction of TRAP-labeled cells with c-fos- and TRAP-double labeling in M1. Bars denote mean and circles individual mice. Error bars, SEM. Control: 20.00% ± 3.45%, n = 17 mice; Early TRAP Trained: 20.65% ± 1.92%, n = 11 mice; Late TRAP Trained: 29.45% ± 4.84%, n = 13 mice. p = 0.235, One-way ANOVA. (H) Fraction of TRAP-labeled cells with c-fos- and TRAP-double labeling in M1, comparing control mice and late TRAP non-learner and learner mice. Bars denote mean and circles individual mice. Error bars, SEM. Control: 20.00% ± 3.45%, n = 17 mice; Late TRAP Non-Learner: 19.75% ± 33.79%, n = 7 mice; Late TRAP Learner: 40.77% ± 7.36%, n = 6 mice. p = 0.999, Control vs. Non-Learner; p = 0.013, Control vs. Learner; p=0.034, Non-Learner vs. Learner, One-way ANOVA with Tukey’s multiple comparison test. (I) Correlation between fraction of c-fos- and TRAP-double labeled cells in M1 and reaching performance (success rate) for individual Early TRAP or Late TRAP mice. Line represents linear regression. Early TRAP mice: n = 11, r^2^ = 0.031, p = 0.607; Late TRAP mice: n = 13, r^2^ = 0.336, p = 0.038, Pearson correlation. *p < 0.05; ns, non-significant

Using this approach, we found that the number of cells activated during early or late stages of learning, either measured by the number of cells expressing tdTomato or by the number of cells that are c-fos positive, did not differ from control mice (Figures 1E and 1F). We next asked if neurons were reactivated to the same degree during early or late training stages. Overall, the fraction of cells that were initially TRAPed (tdTomato-positive) and subsequently reactivated (c-fos immunostaining positive) was not different between control, early, and late TRAP groups (Figure 1G). However, we noticed that within the late TRAP group, mice that learned the reaching task well had a significantly higher level of reactivation compared to non-learners (Figure 1B and 1H). Remarkably, when comparing the level of neuron reactivation with the rate of successful reaches on the days of TRAPing, we found a strong correlation in late TRAP mice, but not early TRAP mice (Figure 1I). These results suggest that mice that learned to perform the reaching task proficiently engage a more defined population of M1 neurons that are reactivated during subsequent execution of the task. On the other hand, mice that have not learned to perform the task well – either during early stages of training or non-learners at late training stages (Figure 1B) – engage a more variable set of neurons that is reactivated to a lower degree.

As a control, we also measured neuronal activation and reactivation during the forelimb reaching task in the primary somatosensory cortex (S1, Figure S1). Here we observed an increased number of c-fos positive cells as well as a higher degree of reactivation in the early TRAP mice compared to controls, while late TRAP mice showed similar levels of activation and reactivation as control mice (Figure S1B-D). In S1, the degree of reactivation did not correlate with the behavioral performance in early or late TRAP mice (Figure S1E and S1F), suggesting that while early training might engage the somatosensory cortex, reactivation of behavior-relevant neurons is specific to M1. Together, these results indicate that by using the c-fos TRAP system, we can identify and genetically label motor engram neurons in M1 that are specifically activated during learning and reactivated during motor memory retrieval in mice that have learned the reaching task.

### M1 dendritic spine plasticity is specific to motor engram neurons

It is well established that motor learning is accompanied by highly coordinated synaptic plasticity in M1, including the rewiring of neuronal connections through the formation and elimination of synapses. Most noticeably, motor learning induces spine reorganization at apical dendrites of L5PNs, where an early increase in spine formation and delayed increase in spine elimination causes a transient increase of spine density during motor training (Xu et al., 2009). Such synaptic plasticity is usually not uniformly distributed across all M1 layer 5 neurons but is only seen in a subset of these neurons during learning (Yang et al., 2014). However, the identity of these dynamic neurons remains elusive. To examine whether neurons that are part of a motor engram are also the neurons that undergo synaptic plasticity during learning, we combined the c-fos TRAP approach with two-photon in vivo imaging of dendritic spines.

We crossed Thy1-YFP mice that sparsely express YFP in layer 5 neurons with the c-fosCre^ER^;Ai9 (c-fos TRAP-tdTomato) mice and implanted a cranial window over their M1. We then repeatedly imaged their apical dendrites daily for four baseline days and subsequently after each forelimb reaching training session for eight days (Figure 2A). Since injecting 4-OHT during late training stages labeled motor behavior-relevant engram neurons in mice learning the forelimb reaching task (Figure 1), we focused on these late TRAP learner mice and injected 4-OHT on training days 7 and 8. A week after daily training, mice were imaged again to acquire green (YFP) and red (tdTomato) channel images to identify which Thy1-YFP dendrites belong to TRAPed engram neurons (Figure 2B). This *post hoc* assignment of dendrite identity allowed us to compare spine dynamics on TRAP dendrites with non-TRAP dendrites in the same mice during motor learning (Figure 2C). To exclude neurons that already expressed tdTomato before 4-OHT injection due to promoter leakage, we took a separate red channel image before TRAPing. We quantified spine density on tdTomato-positive TRAP dendrites as well as nearby tdTomato-negative non-TRAP control dendrites in trained mice and control mice that were only exposed to the training chamber without training (Figure 2A). To verify that our results were not caused by different YFP expression levels or two-photon image clarity between TRAP and non-TRAP dendrites, we quantified baseline fluorescence intensity of the imaged dendrites, which was comparable between groups (Figure S2A). We also detected no difference in baseline spine density between TRAP and non-TRAP dendrites from trained and control mice (Figure S2B). Comparing the overall average spine density between control and trained mice during motor learning, we saw a trend towards a higher density in trained mice, as has been reported before (Xu et al., 2009), though it did not reach statistical significance (Figure 2D and S2C). However, comparing only TRAP dendrites of control and trained mice, revealed a significantly higher spine density in trained mice compared to control mice (Figure 2E and S2D). This difference emerged because TRAP dendrites in trained mice exhibited a transient increase in spine density (Figure 2F and S2E), which was not observed in non-TRAP dendrites of trained mice or either TRAP or non-TRAP dendrites of control mice (Figure 2G and S2F). This transient spine density increase in trained TRAP dendrites is likely caused by an increased level of spine formation and a delayed increase in spine elimination (Xu et al., 2009).

**Figure 2:**
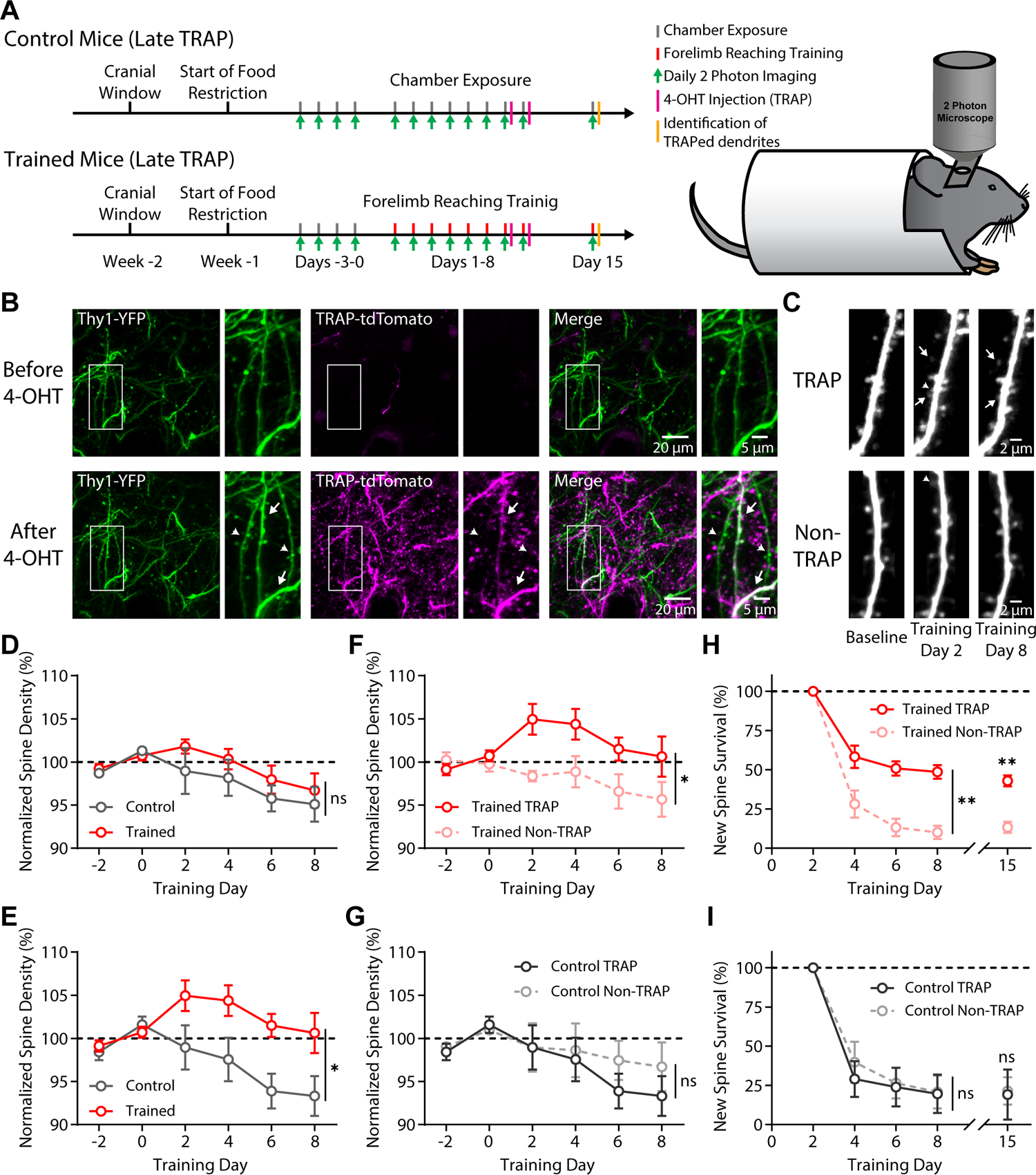
Dendritic spine plasticity during motor learning is specific to engram neurons in motor cortex. (A) Experimental timeline and schematic drawing of in vivo two-photon imaging experiment. (B) Representative image showing the post-hoc identification of dendrites from TRAP neurons. Arrow heads denote dendrites from non-TRAP neurons (Thy1-YFP signal only, green), arrows denote dendrites from TRAP neurons (overlap of Thy1-YFP signal (green) and TRAP-tdTomato signal (magenta), white). (C) Representative images of dendrites from TRAP (top) and non-TRAP (bottom) neurons repeatedly imaged throughout training. Arrow heads denote spines formed during first two days of training that did not persist, arrows denote spines formed during first two days of training and persisted until end of training. (D) Average spine density normalized to baseline spine density in trained (red) and control (gray) mice over the course of motor learning. Control: n = 5 mice with 22 dendritic segments and 1019 spines; Trained: 4 mice with 20 dendritic segments and 866 spines. p = 0.317, 2-way repeated-measures ANOVA. (E) Average normalized spine density of TRAP dendrites in trained (red) and control (gray) mice. Control TRAP: n = 5 mice with 11 dendritic segments and 470 spines; Trained TRAP: n = 4 mice with 10 dendritic segments and 436 spines. p = 0.040, 2-way repeated-measures ANOVA. (F) Average normalized spine density of TRAP dendrites (red solid line) and non-TRAP dendrites (light red dashed line) in trained mice. TRAP: n = 4 mice with 10 dendritic segments and 436 spines; Non-TRAP: 4 mice with 10 dendritic segments and 430 spines. p = 0.044, 2-way repeated-measures ANOVA. (G) Average normalized spine density of TRAP dendrites (gray solid line) and non-TRAP dendrites (light gray dashed line) in control mice. TRAP: n = 5 mice with 11 dendritic segments and 470 spines; Non-TRAP: 5 mice with 11 dendritic segments and 549 spines. p = 0.521, 2-way repeated-measures ANOVA (H) Survival of learning-induced spines (formed on first two days of training) on TRAP dendrites (red solid line) and non-TRAP dendrites (light red dashed line) in trained mice. TRAP: n = 4 mice with 36 newly formed spines; Non-TRAP: 5 mice with 27 newly formed spines. p = 0.008, 2-way repeated-measures ANOVA. Difference persisted at day 15. TRAP: 43.0% ± 3.5%; Non-TRAP: 13.4% ± 3.6%. p = 0.001, paired t-test. (I) Survival of spines formed on first two days on TRAP dendrites (gray solid line) and non-TRAP dendrites (light gray dashed line) in control mice. TRAP: n = 5 mice with 25 newly formed spines, Non-TRAP: 5 mice with 29 newly formed spines. p = 0.766, 2-way repeated-measures ANOVA. No difference at day 15. TRAP: 19.2% ± 16.0%; Non-TRAP: 21.5% ± 8.8%. p = 0.909, paired t-test. *p < 0.05; **p < 0.01; ns, non-significant.

Newly formed spines are also selectively stabilized during learning, where increased levels of new spine survival correlate with behavioral performance (Albarran et al., 2021). Therefore, we assessed the survival of spines that were newly formed during the first two days of training and found that spine survival is indeed significantly higher in TRAP dendrites of trained mice compared to non-TRAP dendrites, a difference that persisted for at least a week after training (Figure 2H). By contrast, survival rates of newly formed spines in control mice were low in both TRAP and non-TRAP dendrites (Figure 2I). Thus, our in vivo imaging data demonstrate that M1 spine plasticity during motor learning preferentially occurs on dendrites belonging to TRAPed engram neurons that are activated in well-trained mice during motor reaching tasks.

### Motor learning strengthens behavior-relevant projections to striatal SPNs

A major output target of M1 neurons is the DLS, which plays a crucial role in generating and organizing voluntary movement. Some studies suggest that striatal activity passively follows cortical activity (Peters et al., 2021), whereas others have found that striatal SPN plasticity is required for motor behaviors (Dang et al., 2006; Santos et al., 2015). This raises the question whether changes in inputs to M1 engram neurons also lead to plasticity at their outputs in the DLS during motor learning.

To study how motor learning modulates corticostriatal connectivity, we selectively expressed Channelrhodopsin 2 (ChR2) in M1 TRAP neurons by injecting AAV5-DIO-ChR2-mCherry into the motor cortex of c-fos-CreER mice (Figure 3A) and administering 4-OHT at early or late stages of training or in control mice (Figure 1A). We then assessed the functional connectivity of M1 engram neurons onto striatal SPNs via whole-cell patch-clamp recording of SPNs with optogenetic stimulation of M1 projections (Figure 3A). Expression of ChR2-mCherry in M1 TRAP neurons and their axon terminals in the DLS was confirmed at 8 weeks after TRAPing (Figure S3A). By measuring the optogenetically evoked excitatory postsynaptic currents (oEPSCs) while stimulating TRAPed M1 neurons, we saw a significantly higher chance to detect a response in recorded SPNs from late TRAP mice (96%), compared to ∼70% in SPNs from control and early TRAP mice (Figure 3B). The oEPSC amplitude in the late TRAP group was also significantly larger than in the control or early TRAP group (Figure 3C and 3D), indicating that motor learning caused an increase in the output of behavior-relevant M1 projections to the striatum. Furthermore, we found that the oEPSC amplitude was similar in mice using either the contralateral or ipsilateral paw for reaching (relative to injection and recording hemisphere), in both early TRAP and late TRAP mice (Figure S3B). These results suggest that learning-induced adaptations in the corticostriatal circuit occur in both hemispheres.

**Figure 3:**
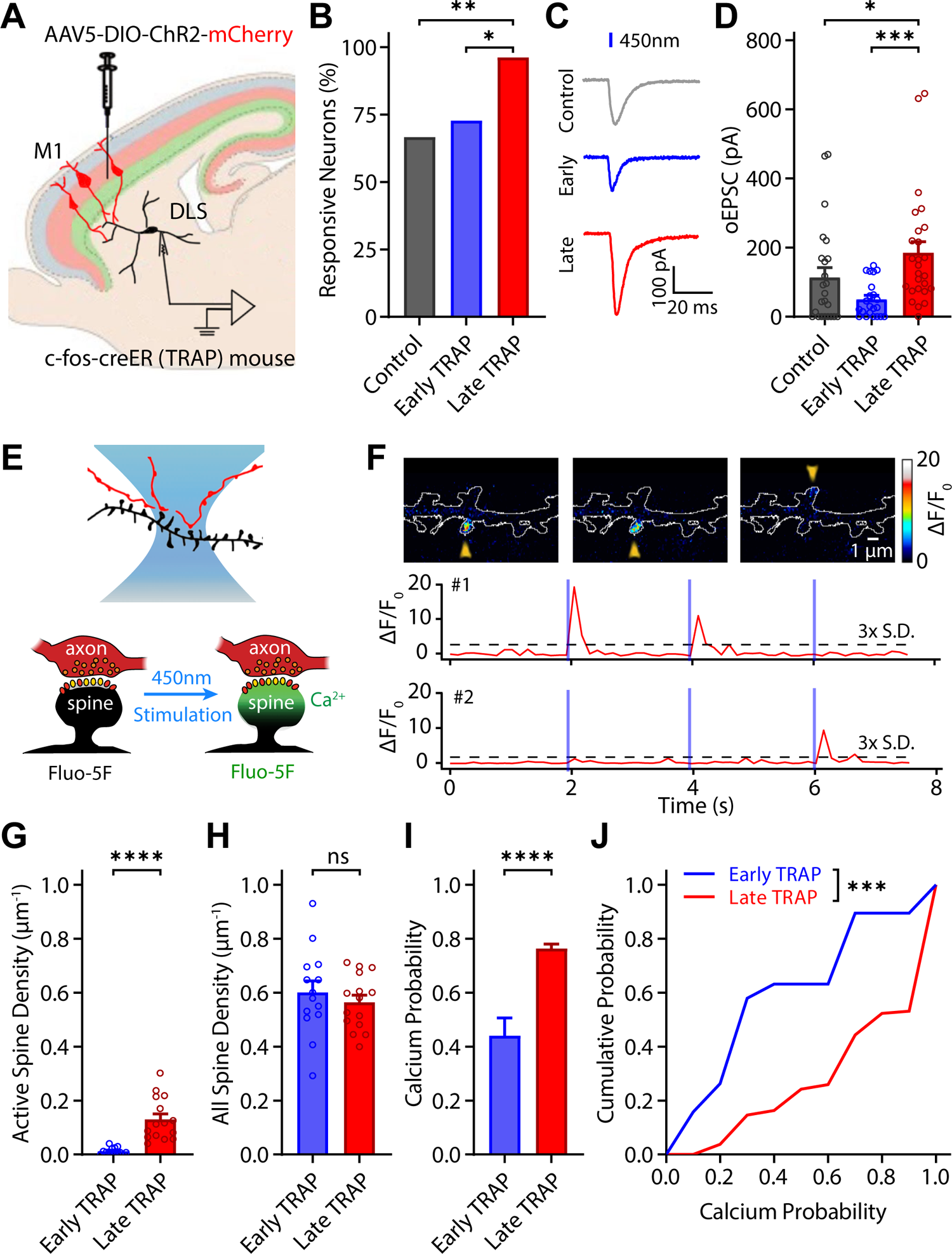
Motor learning strengthens behavior-relevant M1 projections to striatal SPNs. (A) Schematic drawing of M1 TRAP neurons and their projection to striatal SPNs. (B) Fraction of SPNs receiving direct projection from TRAPed M1 neurons. Higher proportion of cells received projections from M1 engram neurons in late TRAP mice compared to early TRAP and control mice. Control: 66.67%, n = 24 neurons from 6 mice; Early TRAP: 72.73%, n = 22 neurons from 7 mice; Late TRAP: 96.15%, n = 26 neurons from 6 mice. Control vs. Late TRAP: p = 0.009; Early vs. Late TRAP: p = 0.038, Fisher’s exact test. (C) Representative traces of optogenetic induced EPSCs (oEPSCs) in striatal SPNs. (D) oEPSCs amplitude in striatal SPNs. oEPSC amplitude in late TRAP mice is higher than in either control or early TRAP mice. Control: 113.27 ± 28.88 pA, n = 24 neurons from 6 mice; Early TRAP: 50.53 ± 11.27 pA, n = 22 neurons from 7 mice; Late TRAP: 184.94 ± 31.97 pA, n = 26 neurons from 6 mice. Control vs Late TRAP: p = 0.034; Early vs Late TRAP: p = 0.001, Kruskal-Wallis test with Dunn’s multiple comparison. (E) Top: Schematic drawing of functional connectome analysis using combined 2-photon Ca^2+^ imaging with optogenetic stimulation. Axonal projections from TRAPed M1 neurons are shown in red and dendrites from striatal SPNs are shown in black. Bottom: Schematic drawing of NMDAR-mediated Ca^2+^ response at a dendritic spine receiving direct input from a TRAP axon terminal during optogenetic stimulation. (F) Top: Representative images of a segment of dendrite during three repeated optogenetic stimulation pulses. Yellow arrowheads indicate Ca^2+^ responses on active spines. Scale bar, 1 µm. Bottom: Fluorescence response from Ca^2+^ imaging using Fluo-5F of the two identified active spines above. Blue bars indicate timing of optogenetic stimulation. (G) Density of active spines on striatal SPNs. SPNs of late TRAP mice receive more projection from M1. Early TRAP: 0.012 ± 0.0034 µm^-1^, n = 15 dendrites from 6 cells and 4 mice; Late TRAP: 0.13 ± 0.021 µm^-1^, n = 10 dendrites from 9 cells and 6 mice. p < 0.0001, Mann-Whitney test. (H) Overall spine density of striatal SPNs. No difference between early and late TRAP mice. Early TRAP: 0.60 ± 0.043 µm^-1^, n = 14 dendrites from 9 cells and 6 mice; Late TRAP: 0.56 ± 0.026 µm^-1^, n = 15 dendrites from 6 cells and 4 mice. p = 0.477, Mann-Whitney test. (I) Ca^2+^ response probability of active spines on SPNs during repeated optogenetic stimulation of TRAPed M1 inputs. Ca^2+^ probability is significantly higher in late TRAP mice than in early TRAP mice. Early TRAP: 0.44 ± 0.066, n = 19 spines from 9 cells and 6 mice; Late TRAP: 0.76 ± 0.017, n = 239 spines from 6 cells and 4 mice, p < 0.0001, Mann-Whitney test. (J) Distribution of Ca^2+^ response probabilities. Distribution in late TRAP mice is shifted toward the higher end. Early vs. Late TRAP: p = 0.001, Kolmogorov-Smirnov test. *p < 0.05; **p < 0.01; ***p < 0.001; ****p < 0.0001; ns, non-significant.

### Motor learning results in an increased number and enhanced efficacy of behavior-relevant corticostriatal synapses on striatal SPNs

We next asked whether the increased M1 motor engram output results in an increased number and/or an increased strength of individual corticostriatal synapses. To address this question, we took a functional connectome approach. Corticostriatal projection neurons form excitatory synapses on the heads of spines along the postsynaptic SPN dendrites (David Smith and Paul Bolam, 1990). We combined optogenetic stimulation of TRAP axons with two-photon Ca^2+^ imaging of postsynaptic SPNs to identify the dendritic spines receiving direct inputs from ChR2-expressing M1 motor engram projections (Figure 3E). Specifically, we performed whole-cell patch-clamp recordings from striatal SPNs with an internal solution containing Alexa594 (50 µM) and the Ca^2+^ indicator Fluo-5F (300 µM), to visualize spine morphology and Ca^2+^ responses, respectively. If a dendritic spine is monosynaptically and functionally innervated by ChR2-expressing M1 axons, upon optogenetic stimulation, we could observe Ca^2+^ transients that are confined to dendritic spines and nearby dendritic segments. A spine was identified as functionally activated by M1 projection neurons (simplified as “active spines”) if its fluorescence intensity exceeded the mean by 3 standard deviations at least once during optogenetic stimulation (Figure 3F and S3C-E).

We compared the number of active spines on SPN dendrites recorded from early and late TRAP mice. Consistent with the results from whole-cell patch-clamp recording data, SPNs from late TRAP mice had a higher density of active spines (Figure 3G), whereas the overall spine density was no different from the early TRAP group (Figure 3H), indicating an increased number of synapses from M1 engram neurons onto SPNs in DLS after motor learning.

We also measured the probability of observing a synaptic response by quantifying the number of postsynaptic Ca^2+^ responses in SPN spines during repeated optogenetic stimulation of ChR2-expressing M1 engram axons (Ca^2+^ probability). This measure can serve as a proxy for the presynaptic release probability and synaptic efficacy. We found that, compared to early TRAP mice, Ca^2+^probability was significantly higher in late TRAP mice, and the distribution across all active spines sampled was shifted towards higher Ca^2+^ probabilities (Figure 3I and 3J). Together, these results indicate that there is an increased number of synaptic connections and an enhanced synaptic efficacy in M1-DLS projections following motor learning.

### Behavior-relevant inputs to SPNs are clustered during motor learning

We next examined the spatial organization of active spines, that are innervated by M1 engram projections, along SPN dendrites because clustering of synaptic inputs has been demonstrated in cortex during motor learning (Fu et al., 2012; Roth et al., 2020) and, in striatal SPNs, co-activation of spatially clustered excitatory synaptic inputs can enhance non-linear synaptic integration and dendritic computation (Du et al., 2017). Taking advantage of the spatial information from Ca^2+^ imaging, we identified and mapped the location of all spines, including active spines, along SPN dendrites (Figure 4A and 4B). In addition, we also used naïve Thy1-ChR2 mice that have not gone through motor learning as a control because, in these mice, ChR2 is expressed in a sparse but random subset of cortical L5PNs (Arenkiel et al., 2007) (Figure 4C). Unlike SPN dendrites in early TRAP mice that have much fewer active spines (≤4 per dendrite), SPN dendrites in late TRAP and Thy1-ChR2 mice have a similar number of active spines (Figure S3F). As a control we also measured the spine morphology of DLS SPNs and found that the distribution of thin, stubby, and mushroom type spines is similar between these groups (Figure S4B). Because the number of active spines was too small in early TRAP group, we focused spatial cluster analysis on the SPN dendrites recorded from late TRAP and Thy1-ChR2 mice.

**Figure 4:**
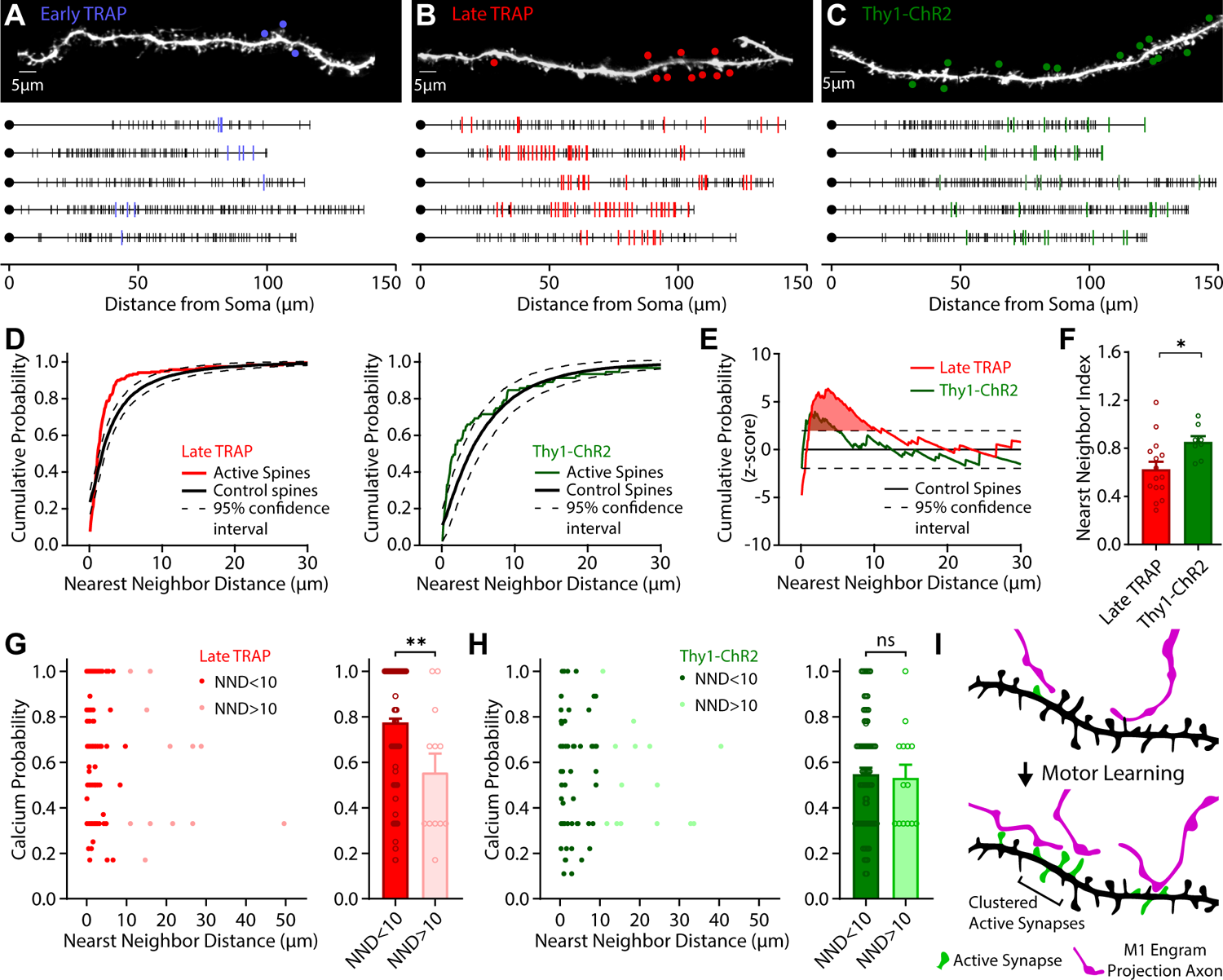
Behavior-relevant inputs to SPNs form clusters during motor learning. (A) Spatial distribution of active spines along SPN dendrites in early TRAP mice. Upper: representative dendrite image with active spines labeled (circles). Lower: 5 representative dendrospinograms, identifying the spatial location of somas (black circles), active spines (colored vertical lines), and non-active spines (black vertical lines). (B) Spatial distribution of active spines along SPN dendrites in late TRAP mice. (C) Spatial distribution of active spines along SPN dendrites in Thy1-ChR2 mice. (D) Cumulative distribution of nearest neighbor distance (NND) for late TRAP (left) and Thy1 mice (right). Red and green lines denote the NND distribution of active spines and black lines denote the distribution from Monte Carlo simulated “control spines” with 95% confidence interval (dashed). (E) Z-score of the NND distribution for late TRAP (red) and Thy1 group (green). Red and green shaded area marks the area under the curve that exceeds the 95% confidence interval of simulated “control spines”, representing the stretch of dendrite with clustered active spines. (F) Nearest neighbor index (NNI) in late TRAP and Thy1 mice. The NNI of late TRAP mice is significantly smaller than that of Thy1-ChR2 mice. Late TRAP: 0.63 ± 0.064, n = 15 dendrites from 6 cells and 4 mice; Thy1-ChR2: 0.85 ± 0.048, n = 8 dendrites from 5 cells and 3 mice. p = 0.013, Mann-Whitney test. (G) Ca^2+^ response probability of synapses inside and outside of clusters in late TRAP mice. Ca2+ probability is higher in spines inside clusters than outside. Left: comparison of NND and spine Ca2+ probability, right: comparison of average Ca^2+^ probability inside (NND < 10 µm) and outside (NND > 10 µm) of clusters. NND < 10: 0.77 ± 0.02, n=227 spines; NND > 10: 0.56 ± 0.02, n = 12 spines. p = 0.007, Mann-Whitney test. (H) Ca^2+^ response probability of synapses inside and outside of clusters in late Thy1-ChR2 mice. No difference between spines inside and outside of clusters. NND < 10: 0.55 ± 0.03, n=77 spines; NND > 10: 0.53 ± 0.02, n = 14 spines. p = 0.836, Mann-Whitney test. (I) Schematic model showing increased numbers of active spines (green) that receive input from M1 engram neurons (magenta) and formation of clusters of active spines on striatal SPN dendrites after motor learning. *p < 0.05; **p < 0.01; ns, non-significant.

To quantify whether active spines are clustered on individual dendrites, we calculated the distance of each active spine to its nearest neighboring active spine and compared this distribution to distances measured in a randomized Monte Carlo simulated “control spine” pool (Figure S4A). We found that out of 15 dendrites we measured in the late TRAP group, 9 dendrites showed significantly smaller nearest neighbor distances among observed active spines compared to randomized control spines, indicating that indeed active spines innervated by M1 engram neurons form clusters along SPN dendrites (Figures 4B). In the Thy1-ChR2 group all 8 dendrites we measured show comparable distances between observed active spines and randomized control spines, indicating that active spines from random set of L5PNs do not cluster on SPN dendrites (Figure 4C). In addition, we plotted the distribution of nearest neighbor distances from all active spines together with that of the simulated “control spines” pool (Figure 4D). We observed that, in late TRAP mice, a large portion of the cumulative distribution curve passed the upper 95% confidence interval of control spine distribution, indicating significant clustering within this nearest neighbor distance range (∼10 µm, Figure 4D). To identify the length of the cluster, the cumulative distribution curves were plotted as a z-score of the simulated randomized pool. Here, we observed significant clustering for distances up to 10 µm in the late TRAP group with a large deviation of the z-score distribution from the upper 95% confidence interval (Figure 4E). Consistently, the nearest neighbor index (NNI), defined as the ratio between nearest neighbor distances of active and randomized spines (Figure S4A), is significantly smaller in the late TRAP group than in the Thy1-ChR2 group (Figure 4F), confirming the formation of synaptic clusters by M1 engram outputs onto postsynaptic SPNs in late TRAP mice.

Since not all active spines were part of clusters in late TRAP trained mice or Thy1 control mice, we asked whether spines within clusters had different presynaptic release probabilities (measured by Ca^2+^ probability) than spines outside of clusters. We, therefore, looked at the relationship between Ca^2+^probability and the clustering of active spines. We found that in the late TRAP group, the Ca^2+^ probability of spines within clusters was significantly higher than that of spines outside of cluster (Figure 4G). Meanwhile, in Thy1-ChR2 mice, there was no difference in Ca^2+^probability between spines within or outside of clusters (Figure 4H). Together, these results indicate that motor learning induces synaptic plasticity that results in strengthening of M1 engram neuron outputs onto postsynaptic striatal SPNs with increased synaptic connectivity, enhanced release probability, and form local clusters along SPN dendrites.

## DISCUSSION

In the present study, we found that motor learning induces the formation of motor engram neurons in M1 that are reliably reactivated during motor memory retrieval in well-trained learner mice. Surprisingly, a well-establish mechanism - spine dynamics on M1 L5PNs during motor learning - was specific to engram neurons in M1 and absent from neighboring non-TRAP neurons, suggesting a potential role of synaptic plasticity in the formation of motor memory engrams. We further found that the outputs of M1 engram neurons also undergo synaptic plasticity during learning, as their projections to SPNs in DLS are strengthened with a higher number of functional connections and a higher presynaptic release probability. Lastly, spines receiving direct inputs from M1 engram neurons are spatially clustered after motor learning, which may further enhance dendritic integration in striatal SPNs.

Reorganization of neuronal activity patterns in M1 and DLS during motor learning has been well documented. It has been shown, for example, that a subset of M1 neurons is modulated during motor reaching (Peters et al., 2014, 2017). The temporal activity pattern of a subset of M1 neurons becomes more defined over the course of motor learning, forming precise and repeatable activity sequences (Peters et al., 2014). Remarkably, the emergence of such neuronal ensembles correlates with the time course in which individual reaching attempts of mice become more precise and stereotyped as well. Similarly, behavior-relevant neurons that we labeled using the genetic TRAP approach become increasingly reactivated with enhanced motor performance of trained mice, suggesting that well-trained mice reactivate the same subset of M1 neurons during individual reaching sessions, whereas naïve or non-learner mice use a different subset of neurons in each session (Figure 1). This enhanced reactivation also coincides with the timeline in which individual reaching attempts become more stereotyped with reduced variability (Albarran et al., 2021). Together, these findings support the concept that neurons with behavior-related activity patterns form a motor memory engram, which becomes more defined with persistent learning.

Synaptic plasticity in the motor cortex plays a crucial role in motor learning. Overall, synaptic inputs to M1 are strengthened in an LTP-like mechanism (Rioult-Pedotti et al., 2000), which on the individual synapse level is driven by the formation, stabilization, and reorganization of synaptic connections (Xu et al., 2009; Yang et al., 2009; Guo et al., 2015; Albarran et al., 2021) as well as through strengthening of existing synapses (Roth et al., 2020). Thus, the question emerges whether these synaptic changes are involved in the development of behavior-related neuronal activity patterns and increased reactivation of engram neurons during motor learning. By showing that spine formation and stabilization are specific to TRAPed engram neurons in M1 (Figure 2), we believe that these synaptic changes play a crucial role in the formation of the memory engram. This is in line with findings in the hippocampus and amygdala, where memory engram neurons possess strengthened synapses and a higher spine density than non-engram neurons (Nonaka et al., 2014; Gouty-Colomer et al., 2015; Ryan et al., 2015; Choi et al., 2018). Enhanced spine formation resulting in higher spine density during learning could result in higher excitability of neurons, which increases the chance of neurons becoming part of an engram. Alternatively, engram neurons could be predetermined to become part of the motor engram because of their intrinsic properties (Yiu et al., 2014; Lisman et al., 2018) and express intracellular molecular mechanisms that enable synaptic plasticity specifically on these neurons. Future experiments directly manipulating synapses on engram neurons (Hayashi-Takagi et al., 2015) would potentially be able to distinguish between these possibilities.

In addition to motor cortex, changes in neuronal activity patterns during motor learning have also been reported in its major output projection target region, the DLS (Jin and Costa, 2010; Barbera et al., 2016; Klaus et al., 2017; Sheng et al., 2019). A longstanding question in the field has been whether the striatum inherits these activity patterns from the cortex or if corticostriatal synapses undergo plasticity to support the reorganization of activity patterns in the DLS. We found that synaptic connections from M1 engram neurons to striatal SPNs are strengthened with a higher number of functional connections (Figure 3), which is likely caused by an increase in the number of synapses from M1 engram neurons since the total number of TRAP neurons in M1 did not change after learning (Figure 1). At the same time, these synapses become strengthened as the increased Ca^2+^ probability at active spines is indicative of an increased presynaptic release probability. Importantly, the plasticity of these corticostriatal projections are from the same M1 engram neurons that undergo dendritic spine plasticity during learning. Together, these results suggest that motor memory traces propagate along M1 engram neurons in the corticostriatal circuit and may be responsible for the formation of striatal neuronal ensembles.

Further supporting the impact of corticostriatal plasticity on DLS activity during learning is our finding that M1 projections to striatal SPNs are spatially clustered (Figure 4). Importantly, since these are inputs from M1 engram neurons this not only shows that these synapses form spatial clusters but also suggests that they are likely activated in a synchronized fashion during a motion sequence since motor behavior-relevant activity in M1 becomes more synchronized after learning (Peters et al., 2014). Such clustering is important for local dendritic computations and can lead to super-linear integration and facilitate the generation of dendritic plateau potentials (Polsky et al., 2004; Losonczy and Magee, 2006; Branco and Häusser, 2011; Stuart and Spruston, 2015; Du et al., 2017).

By defining the plasticity mechanisms of engram neurons in the corticostriatal circuit during motor learning, we further pave the path for studying the cellular and molecular mechanisms involved in corticostriatal plasticity and for understanding how such circuit plasticity and ensemble dynamics are disrupted in movement disorders, such as Parkinson’s disease (Parker et al., 2018) and L-DOPA induced dyskinesia (Girasole et al., 2018). Further characterization of dopaminergic and other inputs to striatal SPNs during motor learning will provide a more complete picture of how cortical and striatal motor engrams are formed and interact during learning.

## STAR METHODS

### Key resources table

| REAGENT or RESOURCE | SOURCE | IDENTIFIER |
| --- | --- | --- |
| **Antibodies** |  |  |
| Anti-c-Fos antibody | Synaptic Systems | Cat# 226 003,<br>RRID:AB_2231974 |
| Donkey anti-Rabbit IgG (H+L) Secondary Antibody, Alexa Fluor 488 | Invitrogen | Cat# A-21206 |
| **Bacterial and virus strains** |  |  |
| AAV5-EF1a-double floxed-hChr2(H134R)-mCherry-WPRE-HGHpA | Addgene | Addgene #20297;<br>RRID:Addgene_20297 |
| **Chemicals, peptides, and recombinant proteins** |  |  |
| 4-Hydroxytamoxifen (4-OHT) | Sigma-Aldrich | Cat# H6278 |
| Fluo-5F, Pentapotassium Salt, cell impermeant | Invitrogen | Cat# F14221 |
| Alexa Fluor 594 Hydrazide | Invitrogen | Cat# A10438 |
| Picrotoxin | Tocris | Cat# 1128 |
| NBQX | Tocris | Cat# 0373 |
| D-Serine | Tocris | Cat# 0226 |
| **Experimental models: Organisms/strains** |  |  |
| Mouse: B6.129(Cg)-Fostm1.1(cre/ERT2)Luo/J (FosCreER) | The Jackson Laboratory | JAX# 021882;<br>RRID:IMSR_JAX:021882 |
| Mouse: Fostm2.1(icre/ERT2)Luo/J (Fos2A-iCreER, TRAP2) | The Jackson Laboratory | JAX# 030323;<br>RRID:IMSR_JAX:030323 |
| Mouse: B6.Cg-Gt(ROSA)26Sortm9(CAG-tdTomato)Hze/J (Ai9) | The Jackson Laboratory | JAX# 007909;<br>RRID:IMSR_JAX:007909 |
| Mouse: B6.Cg-Gt(ROSA)26Sortm14(CAG-tdTomato)Hze/J (Ai14) | The Jackson Laboratory | JAX# 007914;<br>RRID:IMSR_JAX:007914 |
| Mouse: B6.Cg-Tg(Thy1-YFP)HJrs/J (Thy1-YFP-H) | The Jackson Laboratory | JAX# 003782,<br>RRID:IMSR_JAX:003782 |
| Mouse: B6.Cg-Tg(Thy1-COP4/EYFP)18Gfng/J (Thy1-ChR2-YFP) | The Jackson Laboratory | JAX# 007612,<br>RRID:IMSR_JAX:007612 |
| **Software and algorithms** |  |  |
| MATLAB | Mathworks | https://www.mathworks.com;<br>RRID:SCR_001622 |
| ScanImage | Pologruto et al., 2003 | https://vidriotechnologies.com;<br>RRID:SCR_014307 |
| Thorimage | Thorlabs | https://www.thorlabs.com/newgrouppage9.cfm?objectgroup_id=9072 |
| IGOR Pro | WaveMetrics | http://www.wavemetrics.com/products/igorpro/igorpro.htm; RRID:SCR_000325 |
| MapManager | Zhang et al., 2015 | https://mapmanager.net |
| ImageJ | Schneider et al., 2012 | https://imagej.net/; RRID:SCR_003070 |
| Prism 9 | GraphPad Software | http://www.graphpad.com; RRID:SCR_002798 |
| Illustrator | Adobe | https://www.adobe.com/products/illustrator.html; RRID:SCR_010279 |
| Other |  |  |
| Food pellets (20mg) | Bio-Serv | Cat# F0071 |

### Resource availability Lead contact

Further information and requests for resources and reagents should be directed to and will be fulfilled by the Lead Contact: Jun B. Ding.

### Materials availability

This study did not generate new unique reagents.

### Experimental model and subject details

#### Animals

All experiments were performed in accordance with protocols approved by the Stanford University Animal Care and Use Committee in keeping with the National Institutes of Health’s Guide for the Care and Use of Laboratory Animals. All animals were kept on a 12 h:12 h light/dark cycle. For all behavior and in vivo imaging experiments mice were kept at a reversed light cycle room allowing experiments to be performed during the animals’ wake phase. Mice were habituated to the reversed light cycle for at least two weeks before experiments. All behavior and in vivo imaging were carried out at consistent times of day to avoid circadian and sleep-related effects. Both male and female mice were used for all experiments. Behavioral experiments were performed with adult mice at 2-4 months of age. Virus injection experiments were performed in young (P18∼P25) mice to allow for expression prior to behavioral experiments. Mouse lines used in this study include FosCreER (JAX 021882), Fos2A-iCreER (TRAP2, JAX 030323), Ai9 (JAX 007909), Ai14 (JAX 007914), Thy1-YFP-H (JAX 003782), and Thy1-ChR2 (JAX 007612), all obtained from The Jackson Laboratory. Both FosCreER and Fos2A-iCreER were used in the present study. We did not observe any difference in the labeling of M1 cortical neurons, therefore, we combined the data obtained from these two mouse lines. Whenever possible, mice were group-housed over the course of experiments.

## Methods details

### Cranial window surgery

At the age of 2-4 months a custom cut square 3×3 mm cranial window (#1 coverslip glass) was implanted over the primary motor cortex in the left-brain hemisphere as previously described (Roth et al., 2020). Briefly, mice were anesthetized using isoflurane and a craniotomy matching the size of the coverslip was cut using #11 scalpel blades (Fine Science Tools). A coverslip was carefully placed on top of the dura within the craniotomy with mild compression of the brain. The window was centered using stereotactic coordinates 2 mm lateral of bregma. The window was sealed to the skull using dental cement (C&B Metabond, Parkell). At the end of surgery, a custom titanium headplate was attached to the skull to fixate the head to the microscope stage during subsequent imaging. Animals were allowed to recover from surgery for 2 weeks before behavior and imaging experiments.

### Single-pellet reaching task

A single pellet reaching task was performed as previously described (Roth et al., 2020; Albarran et al., 2021). In brief, male and female mice were given restricted access to food to reach and maintain 85% − 90% of their initial body weight. The training chamber consisted of a Plexiglas box with a vertical slit opening as described in (Xu et al., 2009). A platform was placed in front of the opening to position a food pellet (F0071, Bio-Serv) or millet seeds for the mouse to reach. During the week before training onset, all mice were habituated to the training chamber (chamber exposure) and tested for their preferred paw. Therefore, a pile of food pellets was placed on the platform and mice were allowed to perform 10-15 reaches for food pellets. The preferred paw was determined by the more frequently used paw or once mice consecutively used the same paw for 5 reaching attempts. For in vivo imaging experiments, mice had four chamber exposure sessions on the four days directly before training onset to acquire baseline spine images. During these mice were placed in the training chamber for 20 min and a small amount of food was provided inside the chamber. During training, mice were trained daily and each training session consisted of 30 reaches or 20 min. Reaches were classified as successful if the mouse used its preferred paw to grasp the food pellet and bring it directly to its mouth. The success rate was defined as the percentage of successful reaches in all reaches. Early TRAP mice were only trained for two consecutive days and late TRAP mice were trained for eight consecutive days. For c-fos immunostaining experiments, mice in both groups were trained for one additional day 6-7 days after the last day of training. Control mice underwent similar procedures but were only exposed to the training chamber without training and were given a similar amount of food pellets in the chamber as the trained mice retrieved. Mice that failed to reach a minimum success rate of 35% throughout training were classified as non-learners. For *in vivo* imaging and slice optogenetics experiments, only learner mice were analyzed.

### Tamoxifen administration

4-Hydroxytamoxifen (4-OHT, H6278, Sigma-Aldrich) was dissolved at 100 mg/ml in 100% EtOH and stored at −20°C. On the day of injection 4-OHT aliquots were sonicated for 30 - 60 min at 40°C and mixed with sunflower seed oil to a dilution of 10 mg/ml by vortexing. Each mouse received a total of two 50 mg/kg 4-OHT doses which were injected intraperitoneally either after day 1 and day 2 of training (Early TRAP) or after day 7 and day 8 of training (Late TRAP). Control mice for c-fos immunostaining experiments were injected either after the first two or after the last two days. Control mice for *in vivo* imaging experiments followed the late TRAP timeline.

### Slice immunostaining

60 min following the last training session (Day 8 for early TRAP or Day 15 for late TRAP mice), mice were anesthetized with isoflurane and transcardially perfused with phosphate-buffered saline (PBS) and 4% paraformaldehyde (PFA). The brain was dissected out and post-fixed in 4% PFA/PBS for an additional 2 h at room temperature. The brain was transferred to a 30% sucrose/PBS solution for at least 48 h and frozen and sectioned coronally into 50 μm thick slices using a sliding microtome (Leica). Free-floating sections were permeabilized in 0.5% Triton/PBS for 30 min and blocked in 1% BSA, 10% Normal Donkey Serum, and 0.1% Triton for 1h at room temperature. Sections were incubated with primary antibodies (Rabbit anti-c-fos, 1:1000, #226-003, Synaptic Systems) in blocking solution overnight at 4°C and then with secondary antibodies (donkey anti-rabbit-Alexa488, 1:500) in PBS containing 1 % BSA for 2 h at room temperature. Washes after the primary and secondary antibody were done with PBS. Slices from motor cortex and somatosensory cortex were mounted in DAPI containing mounting media (VectaShield H-1500, Vector Laboratories) and multi-channel tiled images were obtained using a laser scanning confocal microscope (LSM900, Zeiss). Representative images shown in figures were median filtered and contrast enhanced.

### In vivo two-photon imaging

Longitudinal *in vivo* two-photon imaging was performed as previously described (Xu et al., 2009; Roth et al., 2020; Albarran et al., 2021) with Thy1-YFP-H mice (JAX 003782) crossed to either FosCreER (JAX 021882) and Ai9 (JAX 007909) or Fos2A-iCreER (TRAP2, JAX 030323) and Ai14 (JAX 007914) mice. Specifically, we repeatedly imaged apical dendrites of L5 pyramidal neurons 10-100 μm below the cortical surface through the cranial window in mice under isoflurane anesthesia. Mice were imaged 1-2h after each training session. Images were acquired using a custom-built two-photon microscope system with a resonant scanner (LotosScan, Suzhou Institute of Biomedical Engineering and Technology) and a 25 x / 1.0 NA water immersion objective lens (Olympus) or a Bergamo II two-photon microscope system with a resonant scanner (Thorlabs) and a 16 x / 0.8 NA water immersion objective lens (Nikon). YFP was excited at 925nm with a mode-locked tunable ultrafast laser (InSightX3, Spectra-Physics) with 15-100 mW of power delivered to the back-aperture of the objective. Image stacks were acquired at 1,024 × 1,024 pixels with a voxel size of 0.12 μm in x and y and a z-step of 1 μm. Imaging frames from resonant scanning were averaged during acquisition to achieve a pixel dwell time equivalent of 1 ns. Up to six imaging regions were acquired for each mouse. For identification of TRAP dendrites that express tdTomato after 4-OHT injection, an additional multi-channel image was taken before 4-OHT injection and 1 week after injection. Therefore, tdTomato was excited at 1100 nm, which also excited YFP to a small degree, allowing registration of images excited at 1100 nm and 925 nm. Representative images shown in figures were created by making z-projections of 3D stacks containing in-focus dendritic segments and were median filtered and contrast enhanced.

### Stereotaxic viral injection

Stereotaxic injections were conducted on (P18∼P25) FosCreER mice (JAX 021882) under isoflurane anesthesia. A volume of 500 nl of concentrated (titer ≥ 1×10^13^ vg/mL) virus (AAV5-EF1a-dflox-hChR2 (H134R)-mCherry-WPRE-HGHpA, #20297, Addgene) was slowly injected in the primary motor cortex of the right hemisphere (AP 1.0 mm, ML 1.5 mm, and DV 1.4 mm from bregma). After injection, the scalp was sutured, and the mice were returned to their housing cages. Animals were allowed to recover from surgery for 2 weeks before behavior experiments.

### Whole-cell slice electrophysiology

Mice (both male and female) were anesthetized and decapitated according to an animal protocol approved by the Stanford University Animal Care and Use Committee. The extracted brains were submerged in ice-cold artificial cerebrospinal fluid (ACSF) containing (in mM) 125 NaCl, 2.5 KCl, 1.25 NaH_2_PO_4_, 25 NaHCO_3_, 15 glucose, 2 CaCl_2_ and 1 MgCl_2_, with continuous bubbling with 95% O_2_ and 5% CO_2_ (300-305 mOsm, pH 7.4). Oblique horizontal slices (300 μm thickness) containing DLS and projections from M1 were prepared as previously described (Ding et al., 2008). Slices were then incubated for 30 min in 34°C ASCF for recovery and then held at room temperature. For patch-clamp recordings, slices were transferred to a recording chamber, held in place with an anchor (Warner Instrument), and mounted onto a microscope (BX51, Olympus) with a customized scanning system for two-photon imaging. The slices were continuously perfused with ACSF at a rate of 2-3 ml/min at 30°C during recording. Patch-clamp recordings were made through a Multiclamp 700B amplifier (Molecular Devices) and monitored with custom-made MATLAB (Mathworks) software. The signal was low-pass filtered at 2.2kHz and digitized at 10kHz (NI PCIe-6259 card, National Instrument). For whole cell voltage-clamp, glass pipettes (2.5-4.5 MΩ) were filled with internal solution containing (in mM) 115 CsMeSO_3_, 10 TEA, 10 HEPES, 5 QX-314 chloride, 4 Mg-ATP, 0.4 Na_3_-GTP, 10 Na_2_-phosphocreatine (280-290 mOsm, adjusted to pH 7.3-7.4 with CsOH). For measuring optogenetic-induced EPSCs, the membrane potential was held at −70 mV and a single optogenetic stimulation pulse was applied to the whole imaging field (10 mW measured at the back-aperture of the objective, 450 nm, 0.5 ms). For all slice electrophysiology experiments, picrotoxin (50 µM) and D-serine (10 µM) were added into ACSF perfusion. Cells with series resistance >25 MΩ were excluded. Since each mouse prefers to use either the left or right paw for reaching, yet the virus injection in M1 and patch-clamp recordings in DLS were consistently performed in the right hemisphere, our stimulation was restricted to M1 projections that were either contralateral or ipsilateral to the preferred paw. We, thus, separated our trained mice into contralateral and ipsilateral subgroups (Figure S3B).

### *Ex vivo* brain slice two-photon imaging

To identify dendritic spines receiving direct input from TRAPed M1 neurons, we used a combination of two-photon Ca^2+^ imaging and optogenetic stimulation similar to previous studies (Little and Carter, 2012; MacAskill et al., 2012; Gökçe et al., 2016). Fifteen minutes after forming whole-cell recording configuration, membrane potential was held at −10 mV to allow for the removal of the Mg^2+^ block of NMDA receptors. In addition to picrotoxin (50 µM) and D-serine (10 µM), NBQX (10 µM) was added to ACSF perfusion to ensure that Ca^2+^ transients were from monosynaptic glutamate transmission. Images were acquired with a custom-built laser scanning microscope controlled with ScanImage (Pologruto et al., 2003) and equipped with a mode-locked tunable Ti:sapphire laser (Mai Tai, Spectra-Physics) and a 60x / 1.1 NA water immersion objective lens (Olympus). Both Alexa 594 and Fluo-5F were excited at a wavelength of 830 nm. For each imaging field, 3 optogenetic stimulation pulses (1 mW, 450 nm, 0.5 ms with interstimulus interval (ISI) of 2050 ms) was applied (Opto Engine LLC) during continuous two-photon morphology and Ca^2+^ imaging with a frame scan rate of 8 Hz and a pixel dwell time of 4 ns. Frame size was 256 x 128 pixels with a pixel size of 0.08 μm in x and y (Galvo-Galvo scanner (Cambridge Technology)). Multiple field-of-view tiles were imaged to cover the entire dendritic segment. Tiles were acquired with at least 33% overlap to allow for offline image stitching using the “Image Stitching” plugin in ImageJ (Preibisch et al., 2009). As a result of tiling, some individual spines were repeatedly imaged and overall each spine was stimulated 9 - 12 times to calculate the Ca^2+^response probability.

## QUANTIFICATION AND STATISTICAL ANALYSIS

### Analysis of slice immunostaining images

Confocal images with fluorescence signal from TRAP-tdTomato and c-fos immunostained cells were analyzed using a semi-automated approach to quantify the number of TRAP and c-fos positive cells in M1 and S1, as well as their overlap. An ROI (∼1.13 mm^2^) covering M1 was defined in both hemispheres in two brain sections at 300 µm and 600 µm anterior to bregma. TRAP and c-fos fluorescence channels were manually thresholded in each ROI and cells were automatically identified and counted using the “Analyze Particles” function in ImageJ after reducing noise pixels (“Despeckle”) and segmenting merged cells (“Watershed”). Overlap between TRAP and c-fos fluorescence was calculated as a fraction of cells with TRAP and c-fos signal (“Double+”) in all TRAP cells. Therefore, the number of pixels with c-fos signal was calculated for each TRAP-positive cell and a cell was defined as double-positive if over 40% of TRAP cell pixels were c-fos-positive. The same approach was applied to S1 where an ROI (∼1.70 mm^2^) was defined over the barrel cortex in both hemispheres in two brain sections at 600 µm and 900 µm posterior to bregma. All analysis was performed blinded to experimental conditions to avoid bias.

### Analysis of in vivo two-photon spine imaging data

Individual dendritic spines were manually identified on dendritic segments and tracked across imaging sessions using the custom software MapManager (https://mapmanager.net) written in Igor Pro (WaveMetrics) as previously described (Zhang et al., 2015). For annotation, the dendritic shaft was first traced using a modified version of the “Simple Neurite Tracer” plugin in ImageJ. Analyzed dendritic segments had a median 3D length of 112.6 µm (min: 27.5 µm, max 181.1 µm). Subsequently, spines were manually marked along the dendrite in 3D image stacks of all imaging sessions. The spine density was calculated for each dendritic segment and averaged per mouse. For survival analysis, spines were further tracked across days by comparing images from different sessions and connecting persistent spines. Spine survival was quantified as the percentage of spines formed during the first two days of training that remained present on a given future training session. Spine density and spine survival data are presented as the average of values from two adjacent imaging sessions (−3 and −2, −1 and 0, 1 and 2, etc.) to increase clarity.

For measuring the dendritic shaft fluorescence intensity at baseline, an ROI covering the dendritic shaft for a 4 μm stretch centered on the connection point of the spine was defined along the traced dendritic shaft. The average fluorescence intensity of all spine-adjacent dendrite ROIs was then averaged for each dendritic segment.

TRAP dendrites were identified posthoc after completion of all imaging experiments. By comparing two-color images taken before and after 4-OHT injection, individual dendrites could be identified that expressed tdTomato after 4-OHT injection but not before. A similar number of non-TRAP dendritic segments was analyzed in each imaging region and non-TRAP dendrites were chosen to match the fluorescence intensity of TRAP dendrites. All analysis was performed blinded to avoid bias. The experimenter was blind to the mouse training condition and TRAP identity of dendrites during analysis.

### Identification of dendritic spines receiving M1 projections

Individual dendritic spines were manually identified on dendritic segments using the custom software MapManager (https://mapmanager.net) written in Igor Pro (WaveMetrics) as previously described (Zhang et al., 2015). For annotation, the dendritic shaft was first traced using a modified version of the “Simple Neurite Tracer” plugin in ImageJ. Along SPN dendrites, spines receiving direct projection from the TRAPed M1 engram neurons were identified by NMDA receptors-mediated Ca^2+^ signal. At each spine head, a small ROI (∼0.05 µm^2^) was selected to extract Ca^2+^ fluorescence intensity changes over time. Spines were identified as “active spines” if their fluorescence intensity during optogenetic stimulation exceeded the baseline mean by 3 standard deviations (S.D.) at least once. Baseline and S.D. was calculated based on the first 15 imaging frames before stimulation. The spatial distribution of active spines and non-active spines along dendrites were then presented as dendrospinograms (Figure 4A-C). Ca^2+^response probability was calculated based on the number of times active spines responded to optogenetic stimulation. Each spine was stimulated for 9 - 12 times total.

### Dendritic spine cluster analysis

To quantify the clustering of identified dendritic spines, the distance of each active spine to its nearest neighboring active spine (nearest neighbor distance, NND) was calculated for each dendrite. The same NND was calculated for a pool of randomized “control spines”, in which the positions of active spines were randomly generated by Monte-Carlo simulation using the cumulative distribution function (cdf) of all spines. A small sample hypothesis test was conducted to test whether the averaged NND from active spines on a given dendrite is significantly different from that of 1000 randomized pools of control spines (Figure S4A). Since the number of active spines was small (≤4 per dendrite) in the early TRAP group, this statistical analysis was only applied to the late TRAP and Thy1-ChR2 groups. The NND distribution of all active spines and randomized control spines regardless of dendrite affiliation was used to identify the distances with significant clustering. To further compare the NND distribution between groups, the cumulative distributions were plotted as z-score, which were standardized by the distribution of each group’s randomized control spine pool (Figure 4E). Also, a nearest neighbor index (NNI) was defined as the ratio between nearest neighbor distances of active and randomized control spines for each dendrite, to compare clustering between groups.

### Statistics

Data analysis was performed in Matlab (MathWorks), IGOR Pro 6.0 (WaveMetrics), and ImageJ. Statistical analysis was performed in Prism 9.0 (GraphPad Software). All data is presented as mean ± SEM (standard error of the mean) in figures and statistical tests, statistical significance (p-values), as well as sample sizes noted in figure legends. Statistical thresholds used: ∗ p < 0.05, ∗∗ p < 0.01, ∗∗∗ p < 0.001, ∗∗∗∗ p < 0.0001, ns: non-significant.

## ACKNOWLEDGMENTS

We thank K. Kwon for assistance with image analysis and members of the Ding lab for insightful discussions and thoughtful comments. This work was funded by NINDS/NIH NS091144 (to J.B.D.), the Klingenstein Foundation (to J.B.D.), and the Parkinson’s Foundation Postdoctoral Fellowship (to R.H.R.).

## Author contributions

F-J.H., R.H.R., Y-W.W., and J.B.D. designed experiments. F-J.H. and Y-W.W. performed and analyzed behavioral, slice electrophysiology and slice imaging experiments. R.H.R. performed and analyzed behavioral and in vivo imaging experiments. F-J.H. and R.H.R. performed and analyzed slice immunostaining experiments. Y.S. and Y.L. provided technical assistance. F-J.H., R.H.R., and J.B.D. wrote the manuscript with input from all authors.

---

**Figure S1 (related to Figure 1):**
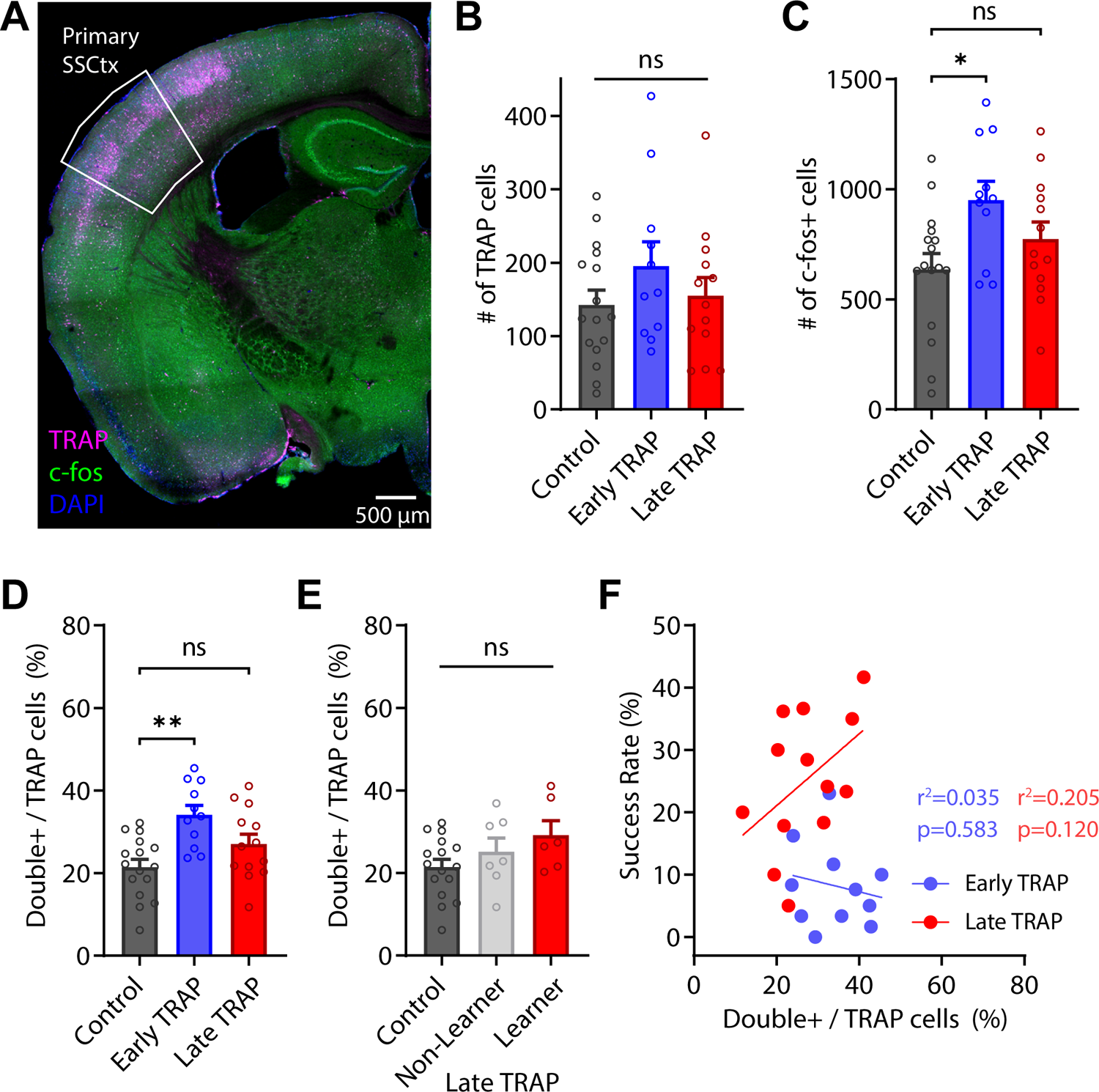
Reactivation of TRAP neurons in somatosensory cortex. (A) Representative image of TRAP-tdTomato (magenta) fluorescence and c-fos immunostaining (green) in primary somatosensory cortex (S1). (B) Number of TRAP-tdTomato labeled cells in S1 (analyzed area: ∼1.70 mm^2^). Bars denote mean and circles individual mice. Error bars, SEM. Control: 142.5 ± 20.02, n = 16 mice; Early TRAP: 195.2 ± 33.33, n = 11 mice; Late TRAP: 154.8 ± 25.04, n = 13 mice. p = 0.342, One-way ANOVA. (C) Number of c-fos-expressing cells in S1. Bars denote mean and circles individual mice. Error bars, SEM. Control: 635.4 ± 72.44, n = 16 mice; Early TRAP: 950.3 ± 85.70, n = 11 mice; Late TRAP: 773.9 ± 77.13, n = 13 mice. Control vs. Early TRAP: p = 0.015; Control vs. Late TRAP: p = 0.340, One-way ANOVA with Tukey’s multiple comparison test. (D) Fraction of TRAP-labeled cells with c-fos- and TRAP-double labeling in S1. Bars denote mean and circles individual mice. Error bars, SEM. Control: 21.46% ± 1.86%, n = 16 mice; Early TRAP: 34.10% ± 2.34%, n = 11 mice; Late TRAP: 27.03% ± 2.38%, n = 13 mice. Control vs. Early TRAP: p = 0.002; Control vs. Late TRAP: p = 0.241, One-way ANOVA with Tukey’s multiple comparison test. (E) Fraction of TRAP-labeled cells with c-fos- and TRAP-double labeling in S1, comparing control mice and late TRAP non-learner and learner mice. Bars denote mean and circles individual mice. Error bars, SEM. Control: 21.46% ± 1.86%, n = 16 mice; Late TRAP Non-Learner: 25.19% ± 3.29%, n = 7 mice; Late TRAP Learner: 29.17% ± 3.531%, n = 6 mice. p = 0.137, One-way ANOVA. (F) Correlation between fraction of c-fos- and TRAP-double labeled cells in S1 and reaching performance (success rate) for individual Early TRAP or Late TRAP mice. Line represents linear regression. Early TRAP mice: n = 11, r^2^ = 0.035, p = 0.583; Late TRAP mice: n = 13, r^2^ = 0.205, p = 0.120, Pearson correlation. *p < 0.05; **p < 0.01; ns, non-significant.

**Figure S2 (related to Figure 2):**
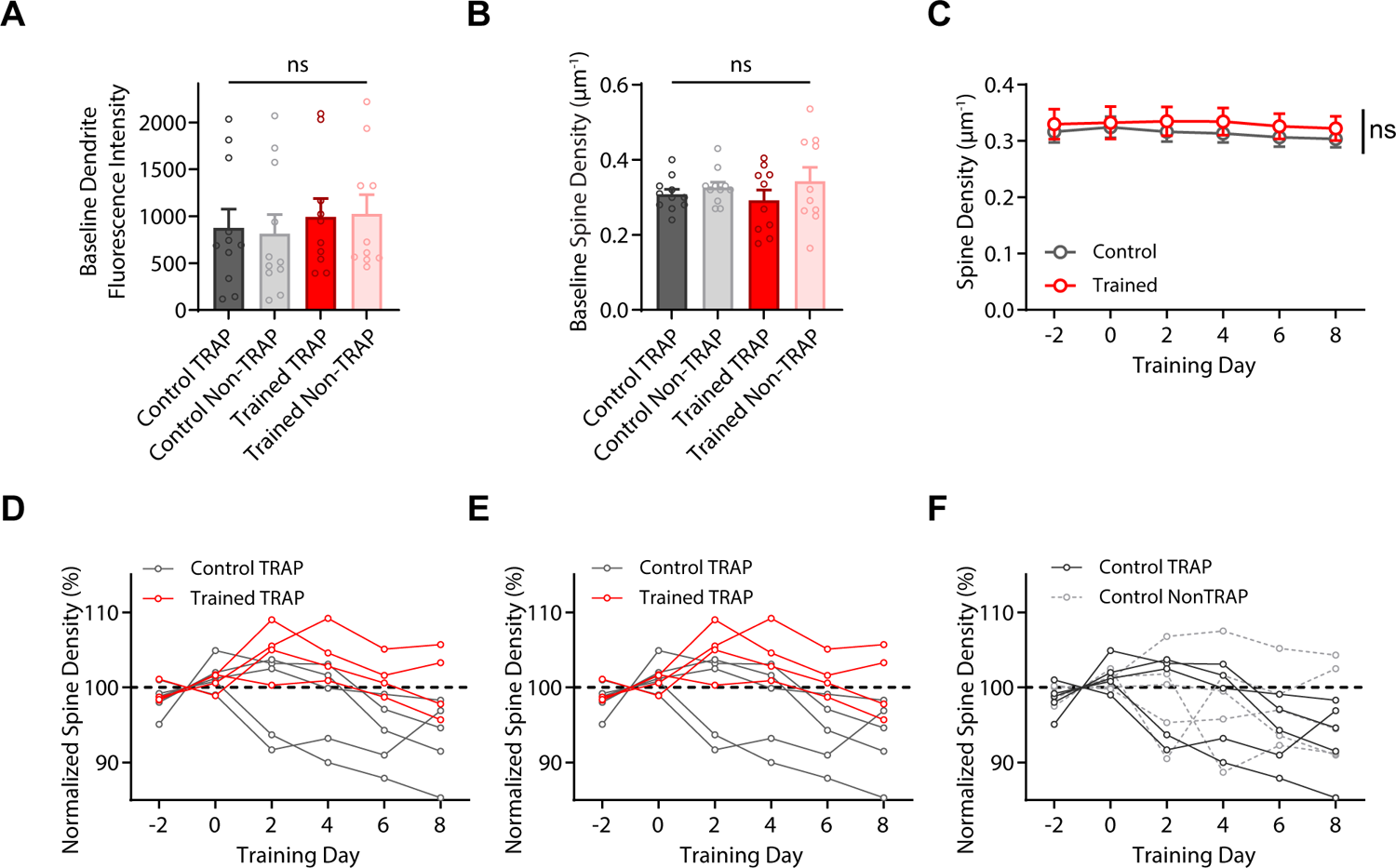
Characterization of spine plasticity during motor learning. (A) Baseline fluorescence intensity of dendritic shaft between TRAP and Non-TRAP segments in trained and control mice shows no difference. Control TRAP: 876.7 ± 198.7, n = 11 dendrites from 5 mice; Control Non-TRAP: 814.6 ± 202.9, n = 11 dendrites from 5 mice; Trained TRAP: 993.7 ± 196.0, n = 10 dendrites from 4 mice; Trained Non-TRAP: 1027 ± 202.5, n 10 dendrites from 4 mice. p = 0.943, Two-way ANOVA, TRAP effect. (B) Baseline fluorescence intensity of dendritic shaft between TRAP and Non-TRAP segments in trained and control mice shows no difference. Control TRAP: 0.308 ± 0.013, n = 11 dendrites from 5 mice; Control Non-TRAP: 0.326 ± 0.014, n = 11 dendrites from 5 mice; Trained TRAP: 0.292 ± 0.027, n = 10 dendrites from 4 mice; Trained Non-TRAP: 0.343 ± 0.037, n 10 dendrites from 4 mice. p = 0.162, Two-way ANOVA, TRAP effect. (C) Average spine in trained (red) and control (gray) mice over the course of motor learning. Control: n = 5 mice with 22 dendritic segments and 1019 spines; Trained: 4 mice with 20 dendritic segments and 866 spines. p = 0.586, 2-way repeated-measures ANOVA. (D) Normalized spine density of TRAP dendrites in individual trained (red) and control (gray) mice. Control TRAP: n = 5 mice with 11 dendritic segments and 470 spines; Trained TRAP: n = 4 mice with 10 dendritic segments and 436 spines. p = 0.040, 2-way repeated-measures ANOVA. (E) Normalized spine density of TRAP dendrites (red solid line) and non-TRAP dendrites (light red dashed line) in individual trained mice. TRAP: n = 4 mice with 10 dendritic segments and 436 spines; Non-TRAP: 4 mice with 10 dendritic segments and 430 spines. p = 0.044, 2-way repeated-measures ANOVA. (F) Normalized spine density of TRAP dendrites (gray solid line) and non-TRAP dendrites (light gray dashed line) in individual control mice. TRAP: n = 5 mice with 11 dendritic segments and 470 spines; Non-TRAP: 5 mice with 11 dendritic segments and 549 spines. p = 0.521, 2-way repeated-measures ANOVA ns, non-significant.

**Figure S3 (related to Figure 3):**
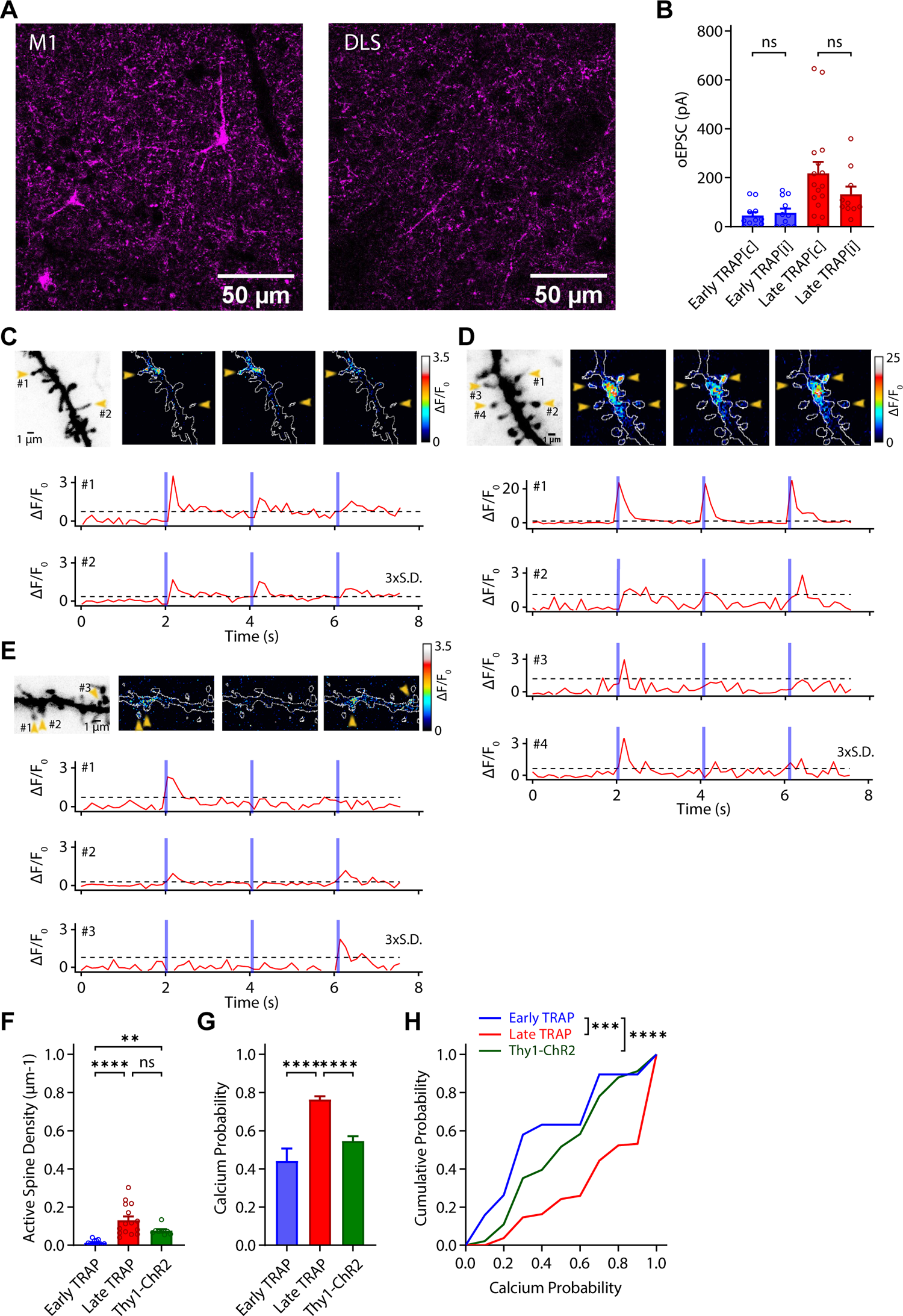
Strengthening of behavior-relevant M1 projection to striatal SPNs during motor learning. (A) Representative confocal images of mCherry fluorescence from c-fos TRAP mice injected with AAV5-DIO-ChR2-mCherry in M1. Injection site in M1 (left) and projections in DLS (right). Scale bar, 50 µm (B) oEPSCs amplitude in striatal SPNs, separated to contralateral and ipsilateral hemispheres based on the preferred paw during reaching. For both early and late TRAP mice, oEPSCs amplitude is similar between the contralateral and ipsilateral mice. Early contralateral: 45.14 ± 14.46, n = 11 from 4 mice; Early ipsilateral: 55.92 ± 17.85 pA, n = 11 from 3 mice; Late contralateral: 218.0 ± 46.91, n = 16 from 4 mice; Late ipsilateral: 132.0 ± 31.77 pA, n = 10 from 2 mice. Early contralateral vs. ipsilateral: p = 0.933; Late contralateral vs. ipsilateral: p = 0.182, Mann-Whitney test. (C) Left: Representative images of a segment of dendrite during three times of repeated optogenetic stimulation. Yellow arrowheads indicate Ca^2+^ responses on active spines. Scale bar, 1 µm. Right: Fluorescence response from Ca^2+^ imaging using Fluo-5F of the identified active spines. Blue bars indicate timing of optogenetic stimulation. (D) Additional examples of active spines, same as (C) (E) Additional examples of active spines, same as (C) (F) Density of active spines on striatal SPNs. SPNs of Thy1-ChR2 mice receive a similar number of projections as late TRAP mice. Both groups have more active spines than early TRAP mice. Early TRAP: 0.012 ± 0.0034 µm^-1^, n = 15 dendrites from 6 cells and 4 mice; Late TRAP: 0.13 ± 0.021 µm^-1^, n = 10 dendrites from 9 cells and 6 mice. Thy1-ChR2: 0.076 ± 0.0087, n = 8 dendrites from 5 cells and 3 mice. Early vs. Late TRAP: p < 0.0001; Early TRAP vs. Thy1-ChR2: p = 0.003; Late TRAP vs. Thy1-ChR2: p > 0.99, Kruskal-Wallis test with Dunn’s multiple comparison. (G) Ca^2+^ response probability of active spines on SPNs during repeated optogenetic stimulation of input projections. Ca^2+^ probability is significantly higher in late TRAP mice than in either early TRAP or Thy1-ChR2 mice. Early TRAP: 0.44 ± 0.066, n = 19 spines from 9 cells and 6 mice; Late TRAP: 0.76 ± 0.017, n = 239 spines from 6 cells and 4 mice; Thy1: 0.55 ± 0.025, n = 91 spines from 5 cells and 3 mice. Early vs. Late TRAP: p < 0.0001; Early TRAP vs. Thy1-ChR2: p = 0.779, Late TRAP vs. Thy1-ChR2: p < 0.0001, Kruskal-Wallis test with Dunn’s multiple comparison. (H) Distribution of Ca^2+^ response probabilities. Distribution in late TRAP mice is shifted toward the higher end. Early vs. Late TRAP: p = 0.001; Late TRAP vs. Thy1-ChR2: p < 0.0001, Kolmogorov-Smirnov test. *p < 0.05; **p < 0.01; ***p < 0.001; ****p < 0.0001; ns, non-significant.

**Figure S4 (related to Figure 4):**
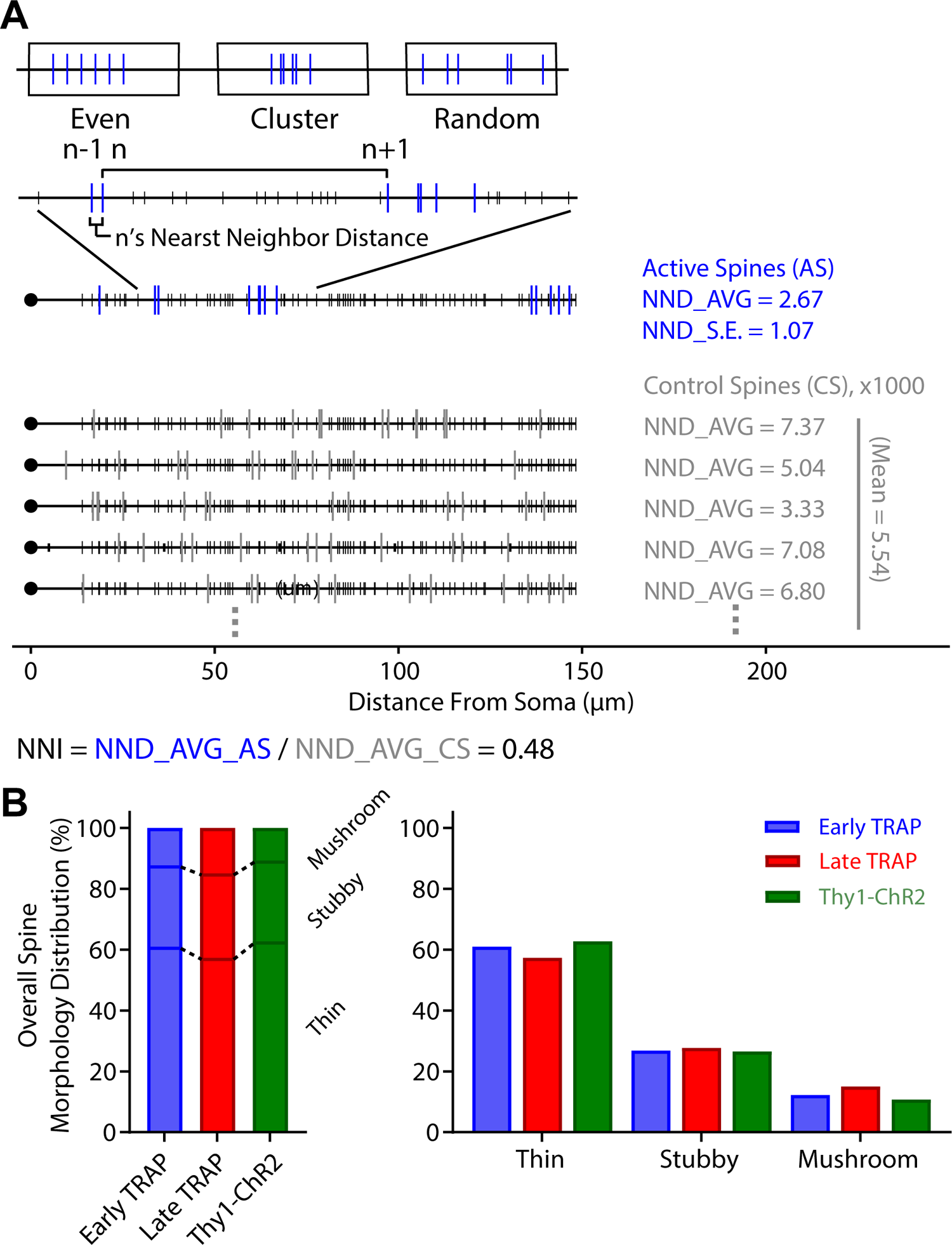
Analysis of SPN spine cluster formation during motor learning. (A) Cluster analysis on representative SPN dendrites. Upper: schematic drawing of evenly, clustered, and randomly distributed spines along a dendrite. Middle: spatial distribution of active spine along a representative SPN dendrite, with the calculated average NND (NND_AVG) and standard error (NND_S.E.). Lower: spatial distribution of Monte-Carlo simulated “control spines” along the representative SPN dendrite, with calculated NND averages. Morphology distribution of all SPN spines shows no significant difference between groups. p = 0.114, Chi-square test.

